# Induction of virus-induced gene silencing and *in planta* validation in cucurbits using the CFMMV-Cm vector

**DOI:** 10.1101/2021.11.18.469169

**Authors:** Sun-Ju Rhee, Yoon Jeong Jang, Jun-Young Park, Gung Pyo Lee

**Author notes:** Author for communication (Gung Pyo Lee). Senior author (Gung Pyo Lee). GL and SR designed the study, SR performed the experiments, analyzed the data, and drafted the manuscript. YJ performed the cloning and histological analysis. JP performed the Agroinoculation and phenotyping. GL supervised the data analysis, revised the manuscript, and funded the project. All authors reviewed and approved the submitted version of the manuscript. The author responsible for distribution of materials integral to the findings presented in this article in accordance with the policy described in the Instructions for Authors (https://academic.oup.com/plphys/pages/general-instructions) is Gung Pyo Lee. **[Email address of Author for Contact]** Sun-Ju Rhee (SR) Yoon Jeong Jang (YJ) Jun-Young Park (JP) Gung Pyo Lee (GL).

## Abstract

Virus-induced gene silencing (VIGS) has been employed for the high-throughput analysis of endogenous gene function. We developed a CaMV 35S promoter-driven cucumber fruit mottle mosaic virus-Cm vector (pCF93) for the efficient generation of viral transcripts in plants. Using the novel pCF93 vector, we identified genes related to male sterility in watermelon (*Citrullus lanatus*), which is recalcitrant to genetic transformation. We previously reported reference-based and *de novo* transcriptomic profiling for the detection of differentially expressed genes between a male fertile line (DAH3615) and its near isogenic male sterile line (DAH3615-MS). Based on the RNA-seq results, we identified 38 *de novo*-exclusive differentially expressed genes (DEDEGs) that are potentially responsible for male sterility. Partial genes of 200∼300bp were cloned into pCF93 which was then inoculated into DAH, a small type of watermelon that enables high-throughput screening with a small cultivation area. In this manner, we simultaneously characterized phenotypes associated with the 38 candidate genes in a common-sized greenhouse. Eight out of the 38 gene-silenced plants produced male sterile flowers with abnormal stamens and no pollens. Gene expression levels in flowers were validated via RT-qPCR. Stamen histological sections from male sterile floral buds and mature flowers showed developmental disruption and shrunken pollen sacs. Based on the current findings, we believe that the novel pCF93 vector and our VIGS system facilitate high-throughput analysis for the study of gene function in watermelons.

**One sentence summary:** The CaMV 35S promoter-driven cucumber fruit mottle mosaic virus-Cm vector (pCF93) facilitates large-scale validation of male sterility-related gene functions in watermelon.

## Introduction

Plant male sterility is defined by flowers producing non-viable pollen or no pollen at all, a failure to dehiscent anthers, and a defective stamen. By avoiding self-pollination, male sterility represents a useful tool for hybrid production, which can increase crop yield by 3.5–200% in various crops (Chen and Liu, 2014; Kim and Zhang, 2018). Thus, insights obtained through functional genomic studies of male sterility can present major benefits for crop breeders and farmers. Within functional genomics, several techniques based on forward and reverse genetics have been developed, including targeting induced local lesions in genomes (TILLING), ethyl methanesulfonate (EMS) mutagenesis, and the production of the transgenic lines via T-DNA tagging and RNA interference. Further, the discovery and introduction of sequence-specific nucleases, such as zinc-finger nucleases (ZFNs) and transcription activator-like effector nucleases (TALENs), has revolutionized gene editing (Gupta and Musunuru, 2014). However, these are difficult to design and are relatively expensive. Over the past three decades, an advanced gene editing technology named ‘CRISPR/Cas9’ system has been developed and introduced for use in humans, animals, as well as plants (Pickar-Oliver and Gersbach, 2019). Although the CRISPR/Cas9 system is a promising tool, it generally requires the generation of transgenic plants via Agrobacterium-mediated or protoplast transformation (Fister et al., 2018; Shan et al., 2020). This bottleneck has been addressed to enable the application of CRISPR/Cas9 in various plants that are recalcitrant to genetic modification. In general, cucurbit plants have extremely low efficiency of transformation. To date, several studies have reported the generation of cucurbit knock-out mutants by using CRISPR/Cas9 (Hu et al., 2017; Tian et al., 2017; Tian et al., 2018). However, the requirement for a stable genetic transformation technique, which is not as laborious and time-consuming, remains.

The virus-induced gene silencing (VIGS) system represents one such alternative as it enables fast and straightforward high-throughput gene screening for functional genomics. Improved VIGS vectors are very recently reported. The Maize dwarf mosaic virus vector could targeted multi endogenous gene simultaneously (Xie et al., 2021) and the Pr CMV-LIC VIGS vector adopted ligation-independent cloning (LIC) strategy for high-throughput cloning (Li et al., 2021). The modified TRV vector is developed for efficient validation system through marker pds gene in front of the multiple cloning site (MCS) (Yamamoto et al., 2021). Efficient and easy-to-handle VIGS systems are in the spotlight for high-throughput screening of the gene functions. VIGS is based on post-transcriptional gene silencing (PTGS), which is a defense mechanism of host cells against viral infection (Robertson, 2004). During plant infection, viral dsRNA is formed through replication. These dsRNAs are degraded by DCL2/DCL4, resulting in the generation of siRNAs. The siRNAs are in turn recruited to endogenous mRNA or viral RNA via the RISC complex based on sequence homology. The targeted mRNA and viral RNA are degraded as a consequence. In order to utilize PTGS in functional genomics, plant virus vectors have been extensively developed in order to apply VIGS systems in various plant species, such as tomato (Liu et al., 2002; Fu et al., 2005; Senthil-Kumar and Mysore, 2011; Rhee et al., 2018), Chinese cabbage (Yu et al., 2019), rice (Purkayastha et al., 2010; Kant et al., 2015), maize (Benavente et al., 2012; Mei et al., 2016), barley (Liu et al., 2016), strawberry (Li et al., 2019), and, more recently, cucurbits (Igarashi et al., 2009; Bu et al., 2019; Liao et al., 2019) as well as banana (Tzean et al., 2019).

We previously constructed a virus vector for functional genomic studies in cucurbits, particularly of their flowers and fruits, using *Cucumber fruit mottle mosaic virus* (CFMMV). CFMMV is a member of the *Tobamovirus* genus and has a monopartite genome of 6.5 kb that is comprised of four ORFs, which encode 127-kDa and 187-kDa proteins involved in virus replication as RNA-dependent RNA polymerases (RdRps), a 30-kDa movement protein (MP) required for cell-to-cell movement of the virus, and a 17-kDa coat protein (CP) that is essential for long-distance movement (Antignus et al., 2001). CFMMV infects a wide range of cucurbits encompassing major commercial crops such as cucumber (*Cucumis sativus)*, melon (*Cucumus melo*), squash (*Cucurbita pepo*), and watermelon. Our groups previously isolated and sequenced CFMMV-Cm from melon to construct a full-length infectious clone (Rhee et al., 2014). We further confirmed that viral *in vitro* transcripts stably replicate in cucurbit fruits. Subsequently, we mapped the sub-genomic protein (SGP) of CP and constructed a CFMMV vector, which successfully expressed EGFP in cucumber, melon, and watermelon (Rhee et al., 2016).

The functional genomics analysis of watermelon has been challenging because of the recalcitrant nature of the species with regard to the generation of transgenic plants. Further, the study of watermelon fruit development demands a relatively long time as well as large areas for cultivation.

In this study, we report an efficient cucurbit-infecting virus vector system based on the CFMMV VIGS vector, which exhibited high and long-lasting gene silencing efficiency in cucurbit plants, with clear phenotypes. Our monopartite CFMMV vector is relatively easy to manipulate and allows the high-throughput screening of gene function in watermelon. This CFMMV VIGS system will help advance the study of gene function in cucurbit reproductive organ development.

## Results

### Constructing an efficient CFMMV vector for VIGS

In order to determine the most efficient VIGS vector, four CFMMV-Cm-based vectors (pCF93, pCF93K, pCF157, and pCF157K) were assessed based on our previous report (Rhee et al., 2016). pCF93 and pCF157 have one 3’NTR, while pCF93 and pCF157 have an additional 3’NTR between the MCS and CP region (Figure 1A). To visualize the gene silencing efficiency of CFMMV-Cm-based vectors, the partial *phytoene desaturase* (*pds*) gene was inserted into each vector. Infection symptoms were observed at 5-6 dpi on the second upper leaf of inoculated leaves, and a photo-bleached phenotype developed at 12-15 dpi (Figure 1B, 1C). The typical photo-bleached phenotype was not observed in *Nicotiana benthamiana* infected with pCF157K-nbpds (Figure 1C). Comparison of the gene silencing phenotypes between pCF93-nbpds and pCF157-nbpds revealed that plants infected with the former had wider white-leaf areas than those of infected with pCF157-nbpds. When the CFMMV-Cm-based vector carrying *PDS* was inoculated into *N. benthamiana*, virus symptoms appeared at 5-6 dpi on the second upper leaf from the inoculated leaves, and photo-bleached phenotypes developed at 12-15 dpi. In *N. benthamiana* infected with pCF157K-nbpds, systemic viral symptoms were observed, while photo-bleached phenotypes did not develop. To confirm the inserted gene stability of the vector, quantitative reverse transcription PCR (RT-qPCR) was conducted using det-F and det-R primers designed based on the flanking regions nearby the MCS. After inoculation with pCF93, pCF93K, and pCF157 carrying nbpds, total RNA was isolated from fourth upper leaves above the inoculated leaves in order to confirm their genomic composition via RT-qPCR. Products of expected size were obtained (blue arrow in Supplemental Figure S2). However, pCF157K-nbpds lost the artificially inserted MCS region and returned to the genome structure of wild-type CFMMV via homologous recombination (Supplemental Figure S2). Although pCF93K-nbpds induced photo-bleaching phenotypes, it exhibited low gene silencing efficiency as compared to pCF93-nbpds and pCF157-nbpds. Thus, the vectors without an additional 3’NTR were better suited for VIGS. We selected pCF93 as the most efficient VIGS vector for gene function analysis.

**Figure 1.**
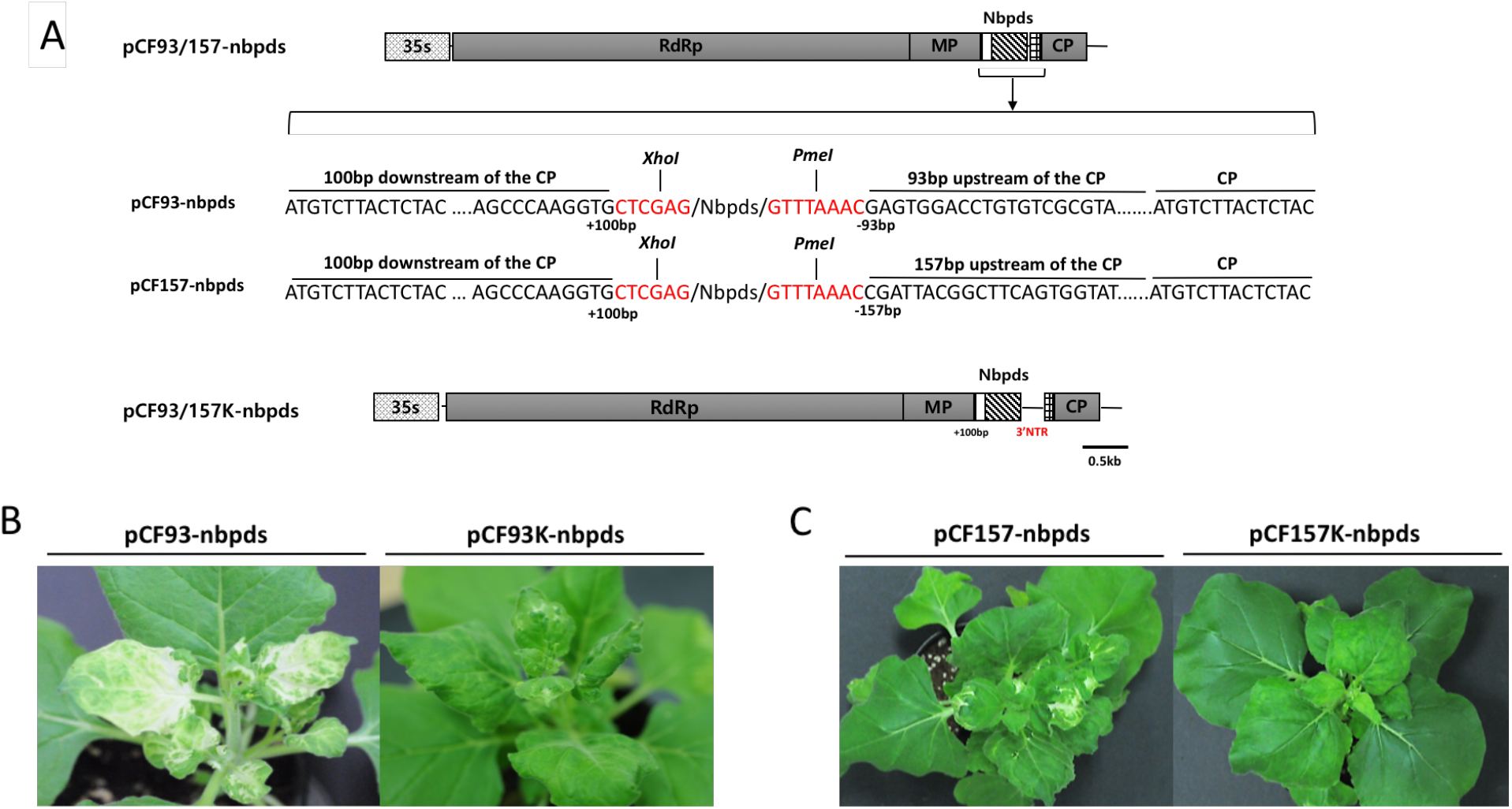
Virus-induced gene silencing of the *PDS* gene resulting in a photobleaching phenotype in *N. benthamiana*. Schematic diagrams of pCF93/157-nbpds and the nucleotide sequence around the MCS region. *Xho*I and *Pme*I are restriction enzyme site for insert cloning. *35S*, Cauliflower mosaic virus 35S promoter; *5’ NTR*, 5’ non translated region; *RdRp*, RNA-dependent RNA polymerase; *MP*, triple gene block; *CP*, coat protein; *3’NTR;* 3’ non translated region (A), comparison of VIGS phenotypes in *N. benthamiana* infected by pCF93-nbpds and pCF93K-nbpds (B) as well as pCF157-nbpds and pCF157K-nbpds (C) at 12 days post-inoculation (dpi). Of the four constructs tested, pCF93 exhibited the most efficient silencing. Thus, we selected pCF93 for VIGS.

### Evaluation of pCF93 as a VIGS vector in cucurbits

*PDS*-carrying pCF93 was inoculated into cotyledons of cucumber, melon, and watermelon in order to validate VIGS efficiency in cucurbits. *N. benthamiana* has been used to evaluate the gene silencing efficiency of various VIGS vectors (Rhee et al., 2016). The *PDS* expression levels of *N. benthamiana* infected with pCF93-nbpds exhibited a 5-fold reduction (Figure 2A). Melon infected with pCF93-cmpds exhibited photobleaching phenotypes in the middle areas of the plant, including the stems and leaves, maintaining these phenotypes in the newly developed leaves until its death (Figure 2B). Likewise, the leaves of watermelon exhibited photobleaching phenotypes and a *PDS* mRNA level reduction of up to 333-fold when compared to p35SCF-Cm_flc_ -infected counterparts (Figure 2C). The cucumber also exhibited *PDS*-silenced phenotypes in the leaves and flowers. The *pds* mRNA level in the leaves of pCF93-cspds-infected cucumbers was reduced about 7-fold relative to that in p35SCF-Cm_flc_ -infected cucumbers. Flowers also exhibited gene silencing phenotypes and an approximately 2.5-fold reduction in *PDS* mRNA expression under pCF93-cspds infection relative to that under p35SCF-Cm_flc_ infection, as determined via RT-PCR (Figure 2D). Of note, all plants infected with the tested virus vectors were transferred from the growth room to a greenhouse under the sunlight in order to enhance *PDS* VIGS efficiency.

**Figure 2.**
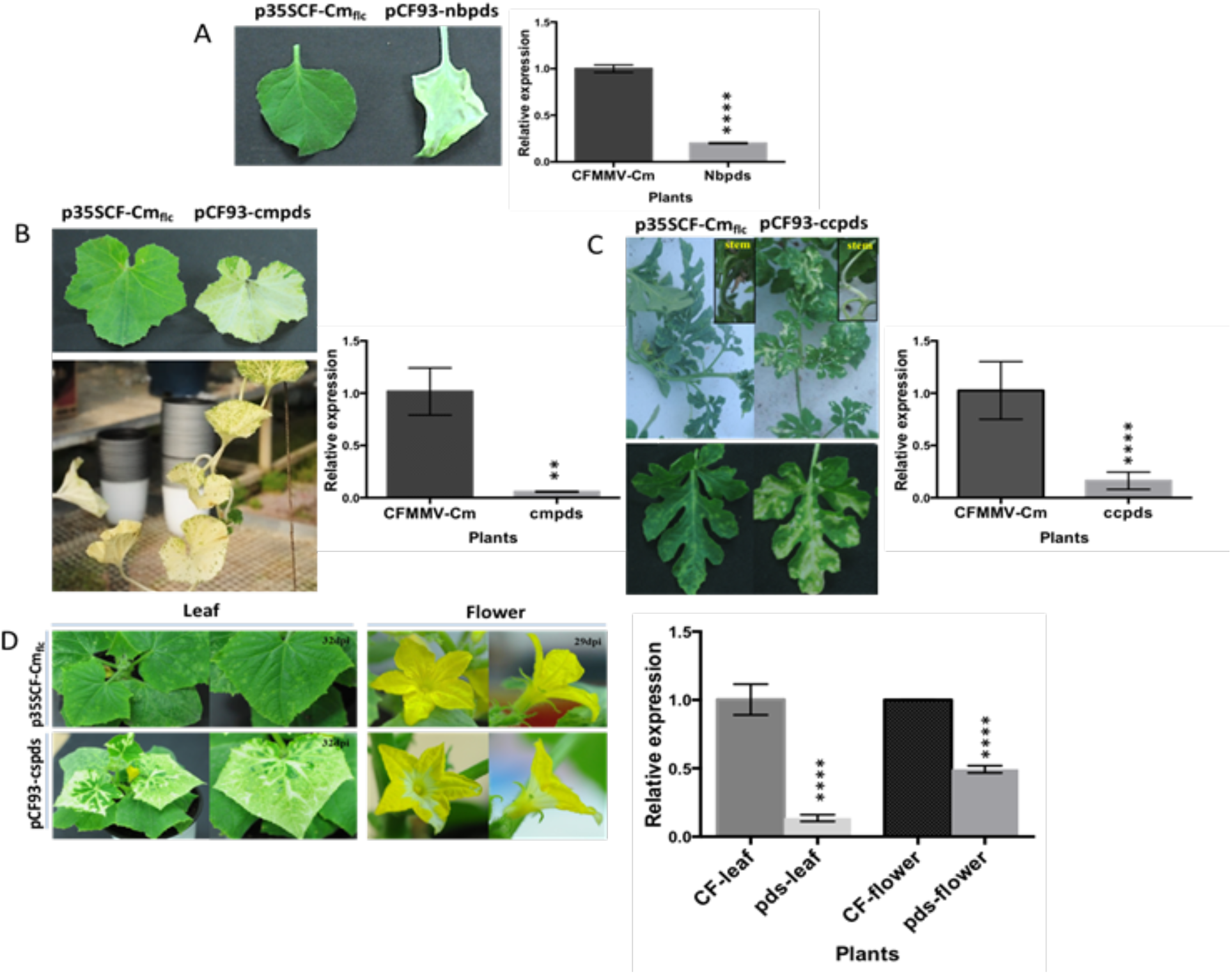
The effects of virus-induced *PDS* silencing on phenotype in various plants. (A) Photobleaching phenotypes and relative expression levels of endogenous *PDS* mRNA in *N. benthamiana* leaves infected with pCF93-nbpds. (B) Comparison of gene silencing efficiency in melon leaves infected with p35SCF-Cm_flc_ (top, left) and pCF93-cmpds (top, right). pCF93-cmpds-induced VIGS phenotypes in melon grown in greenhouse (Bottom). (C) Photobleaching phenotypes on watermelon leaves and stem infected with pCF93-ccpds. Relative expression levels of *PDS* after silencing via pCF93-ccpds compared to those in watermelon infected with wild-type CFMMV, as determined via real-time qPCR. (D) Photobleaching of cucumber leaves and flowers infected with pCF93-cspds. *PDS* mRNA levels in cucumber leaves and flowers infected with p35SCF-Cm_flc_ (denoted as CF-leaf and CF-flower, respectively) and pCF93-cspds (denoted as pds-leaf and pds-flower, respectively). Relative *pds* expression was determined using *PDS*-specific primers and normalized to GAPDH expression. (A) 18S rRNA expression level (B, C, and D). Relative expression levels were determined by setting arbitrary value (1.0) to the expression level of CFMMV-Cm, which used as a calibrator sample. Statistical analysis was performed using the Student’s t-test (****p < 0.0001 and 0.001<**p< 0.01).

### Evaluation of *PDS* gene silencing efficiency in three watermelon cultigens

Flowers and fruits are the major reproductive organs giving rise to edible parts of plants, especially cucurbits. Therefore, many researchers have studied the functional genomics of cucurbit fruit and flower development (Lin et al., 2007; Wang et al., 2010; Manzano et al., 2014; Switzenberg et al., 2015). *pds* carrying pCF93 was Agro-inoculated to the cotyledons of watermelon, and the plants were grown in greenhouse conditions without any additional infections. Three cultigens of watermelon, namely ‘2401’, ‘Chris cross’, and ‘DAH’ (a small-fruit type of watermelon), were used for evaluating pCF93-ccpds VIGS efficiency. Watermelon fruits were harvested at 45 days after pollination (dap) and exhibited photobleaching phenotypes on the peels (Figure 3A). When the watermelon plants were infected with pCF93-ccpds, the fruit peels and flesh turned white. As the red flesh of watermelon is packed with lycopene and β-carotene (Tomes et al., 1963), we presumed that the color change to white was caused by photo-bleaching, which is responsible for the degradation of chlorophyll. White-colored flesh can also develop as a result of the inhibition of lycopene and β-carotene biosynthesis. Thus, lycopene and β-carotene content were estimated via HPLC analysis, and *PDS* expression was determined via semi-RT-qPCR. The lycopene and β-carotene contents of the fruits (45 dap) decreased significantly in all three watermelon cultigens. The flesh color of all tested watermelon fruits became white, which was attributed to the blockade of lycopene synthesis. The lycopene and β-carotene contents of pCF93-ccpds-infected watermelon decreased approximately 3- to 110-fold and 2.7- to 21-fold, respectively, as compared to those in p35SCF-Cm_flc_ -infected fruit (Figure 3B, C, and D). *PDS* gene expression was suppressed in all watermelons subjected to *PDS* silencing (Figure 3E). Chlorophyll a content in the peels of pCF93-ccpds-infected watermelons was reduced about 2.5- to 15-fold compared to that of p35SCF-Cm_flc_ -infected plants, while chlorophyll b content exhibited a 1.4- to 8-fold reduction. Total chlorophyll was reduced about 1.13- to 2.5-fold (Figure 3F and G). Of note, lycopene and β-carotene content were successfully repressed in the fruits of all three watermelon cultigens.

**Figure 3.**
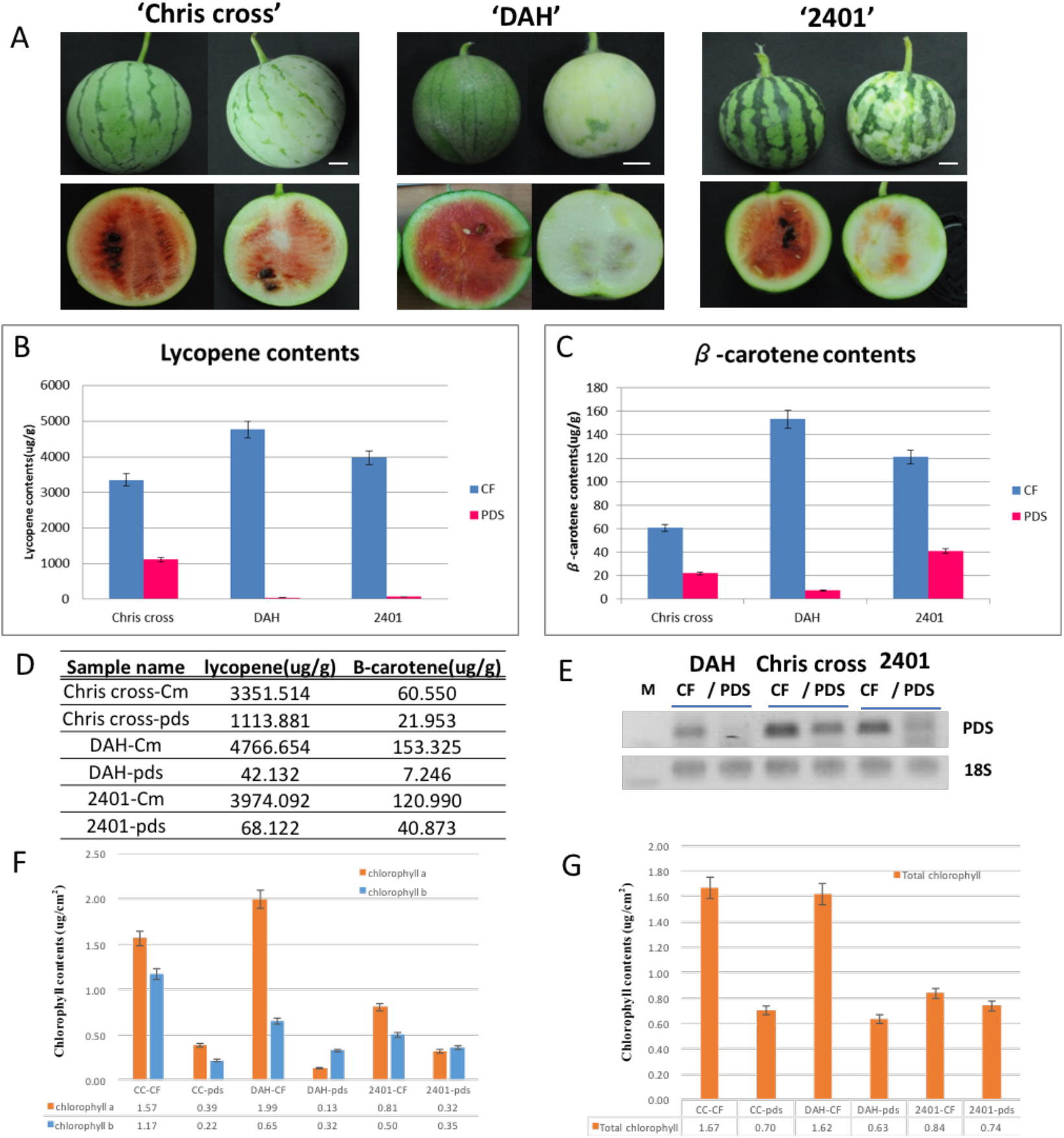
Estimation of lycopene and β-carotene contents in watermelon fruits (‘Chris cross’, ‘DAH’, and ‘2401’) and semi-quantitative RT-qPCR for the detection of endogenous phytoene desaturase (PDS) expression. VIGS phenotypes in the fruits of three cultigens of watermelon. Scale bars=2 cm (A). Quantification of lycopene (B) and β-carotene (C) contents in flesh of watermelon fruits infected with p35SCF-Cm_flc_ (blue bar, indicated as ‘CF’) and pCF93-ccpds (pink bar, indicated as ‘PDS’). Lycopene and β-carotene contents were determined via HPLC. Lycopene and β-carotene contents presented in a table (D), semi-quantitative RT-Qpcr was performed using a ccpds-specific primer set (E). Estimation of Chlorophyll a and b contents in peels of watermelon infected by CFMMV vector (F), Calculated total chlorophyll contents (G). Watermelon cultigens ‘Chris cross’, ‘DAH’ and ‘2401’ infected by p35SCF-Cm_flc_ are denoted as ‘CC-CF’, ‘DAH-CF’ and ‘2401-CF’, respectively. ‘Chris cross’, ‘DAH’ and ‘2401’ infected by pCF93-ccpds are denoted as ‘CC-pds’, ‘DAH-pds’ and ‘2401-pds’, respectively.

### Verification of candidate genes related to male sterility in watermelon by using the CFMMV-VIGS system

We previously conducted reference- and *de novo* RNA-seq to detect differentially expressed genes (DEGs) between the MS and MF lines of DAH3615 (Rhee et al., 2015; Rhee et al., 2017). We identified a total of 38 *de novo*-exclusive DEGs (DEDEGs) and inserted partial genes into pCF93 vectors for gene function analysis (Supplemental Table S1). Watermelon plants infected with vectors for the silencing of each DEG were simultaneously cultured in the greenhouse and studied.

Silencing eight of the DEDEGs, which included calcium-dependent protein kinase 2 (*CDPK2*), elongation factor 1-alpha (*EF1a*), late embryogenesis abundant protein 1 (*LEA1*), LIM domain protein (*LIM*), mitotic spindle assembly checkpoint protein MAD2 (*MAD2*), polygalacturonase (PG), fasciclin-like arabinogalactan protein 5 (*FLA5*), and expansin-A9 (*EXPA9*), resulted in typical male sterility phenotypes (Table 1).

**Table 1.** Eight out of the 38 putative genes from reference-based and *de novo* transcriptome data, which were silenced in watermelons via VIGS.

| Gene symbol | Research | Uniprot_ID | Gene_ID | Annotation |
| --- | --- | --- | --- | --- |
| V4 | Reference-based | Q3YAT0_PETIN | Cla010883 | Calcium-dependent protein kinase 2 |
| V8 | Reference-based | B9SPV9_RICCO | Cla000553 | Elongation factor 1-alpha |
| V9 | Reference-based | LEA1_CICAR | Cla007634 | Late embryogenesis abundant protein 1 |
| V10 | Reference-based | Q306K1_BRANA | Cla001608 | LIM domain protein |
| V12 | Reference-based | D2V3A0_NAEGR | Cla009991 | Mitotic spindle assembly checkpoint protein MAD2 |
| V15 | Reference-based | E3VSV7_CUCPE | Cla016924 | Polygalacturonase |
| V24 | De novo | FLA5_ARATH | c9925_g1 | Fasciclin-like arabinogalactan protein 5 |
| V28 | De novo | EXPA9_ARATH | c29764_g1 | Expansin-A9 |

The coding sequence (CDS) information was obtained from the watermelon genome database of the Cucurbit Genomics Database (http://cucurbitgenomics.org/), and candidate gene primers were designed to detect the insertion of pCF93 (Table S2). All DAH plants infected with the eight constructs produced complete or partially abnormal MS flowers in the same plant (Fig. S3 and Fig. 4).

**Figure 4.**
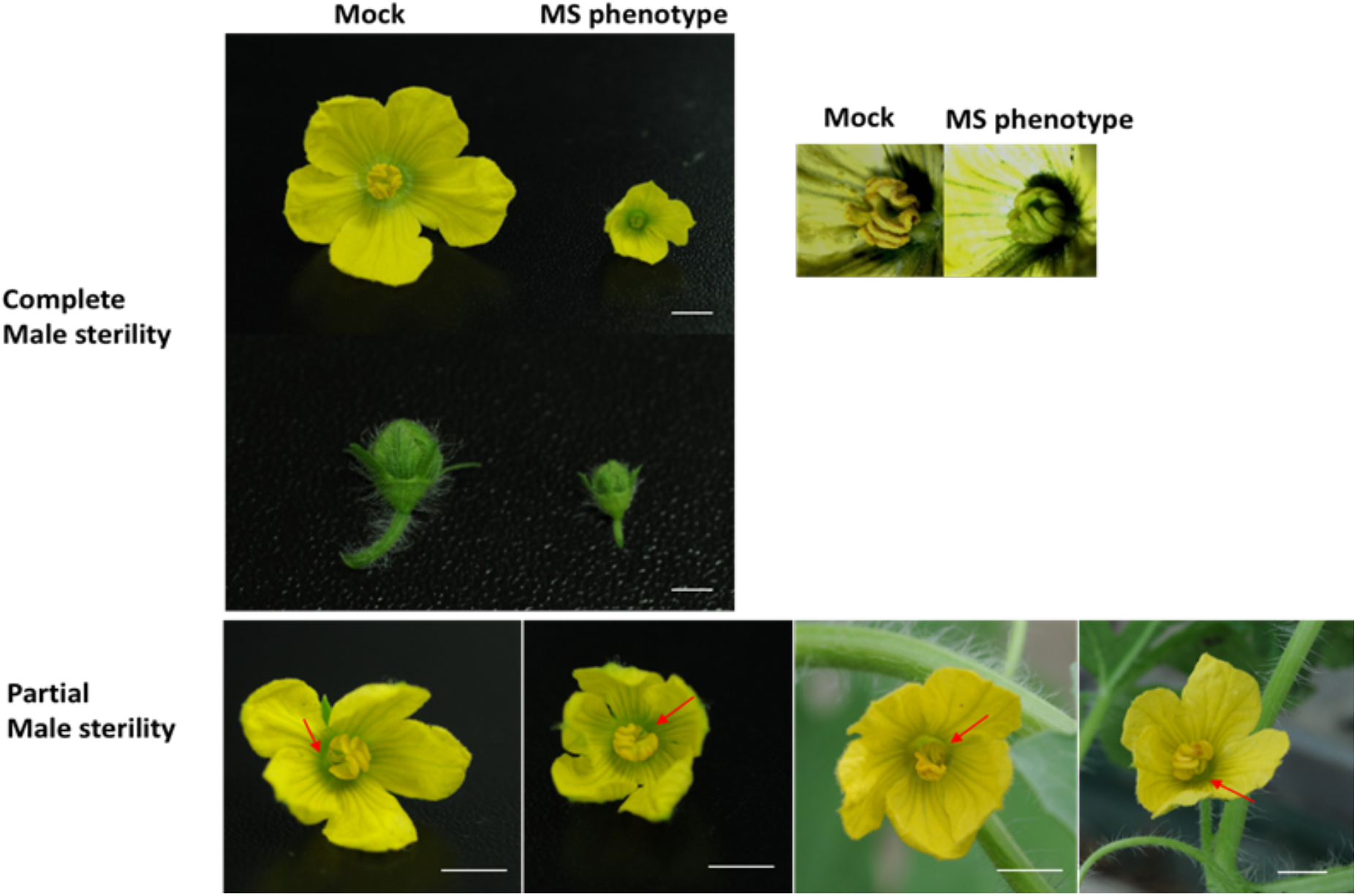
Male sterile flowers developed in candidate gene-silenced watermelon. Complete male sterility observed in candidate gene-silenced plants. Some of the flowers exhibited partial male sterility. Representative flowers in the top panel were collected from pCF93-pds-int (denoted as ‘mock’) and EXPA9-silenced plants (denoted as ‘MS phenotype’). Partially abnormal male sterility flowers developed in LIM-, MAD2-, FAP5, and PG-silenced plants (from left to right, respectively). Red arrows indicate the aborted stamen. Scale bars=0.5 cm.

The wild-type watermelon male flower consists of three individual stamens. The complete MS flower exhibited an almost 3-fold size reduction as well as small green immature stamens and no pollens. The partially abnormal MS flower showed developmental disruption of one or two stamens as well as no pollens. However, the normally developed stamens of partially abnormal MS flowers produced pollens. The *LEA1*-silenced plant produced complete MS flowers but no partially abnormal MS flowers. The seven remaining gene-silenced plants produced both complete and partially abnormal MS flowers. The phenotypes of all virus-induced MS flowers appeared similar to each other as well as to the male sterile line ‘DAH3615-MS’. To characterize anatomical differences between MS and MF stamens, histological cross-sections were evaluated. MF floral buds and flowers were collected from mock-inoculated plants infected with pCF93-pds-int. DAH3615-MS did not produce pollen grains, and stamens appeared immature based on visual examination. MF floral buds and flowers produced normal pollen sacs and pollen, whereas MS showed developmental disruption of tapetum cells and microspore mother cells. The mature flowers had atrophied pollen sacs that consisted of abnormal shrunken cells (Figure 5, indicated by red arrow). RT-qPCR was then performed to determine the expression of the eight DEDEGs and explore their interaction. We collected three mature flowers showing a complete MS phenotype and pooled their total RNA. Most of the genes exhibited a tendency toward suppression in virus-infected MS flowers (Figure 6). The *LEA1*-silenced flowers exhibited dramatic induction of the *LIM* gene, approximately 290-fold compared to that of pds.int-infected plants. Conversely, the *LIM*-silenced flowers exhibited reduced *LEA1* expression. *EXPA9* expression was induced via *FLA5* downregulation, while *CDPK2, MAD2, PG*, and *FLA5* were suppressed in all flowers showing the MS phenotype. Notably, all MS phenotype flowers exhibited extreme reduction in *MAD2* expression.

**Figure 5.**
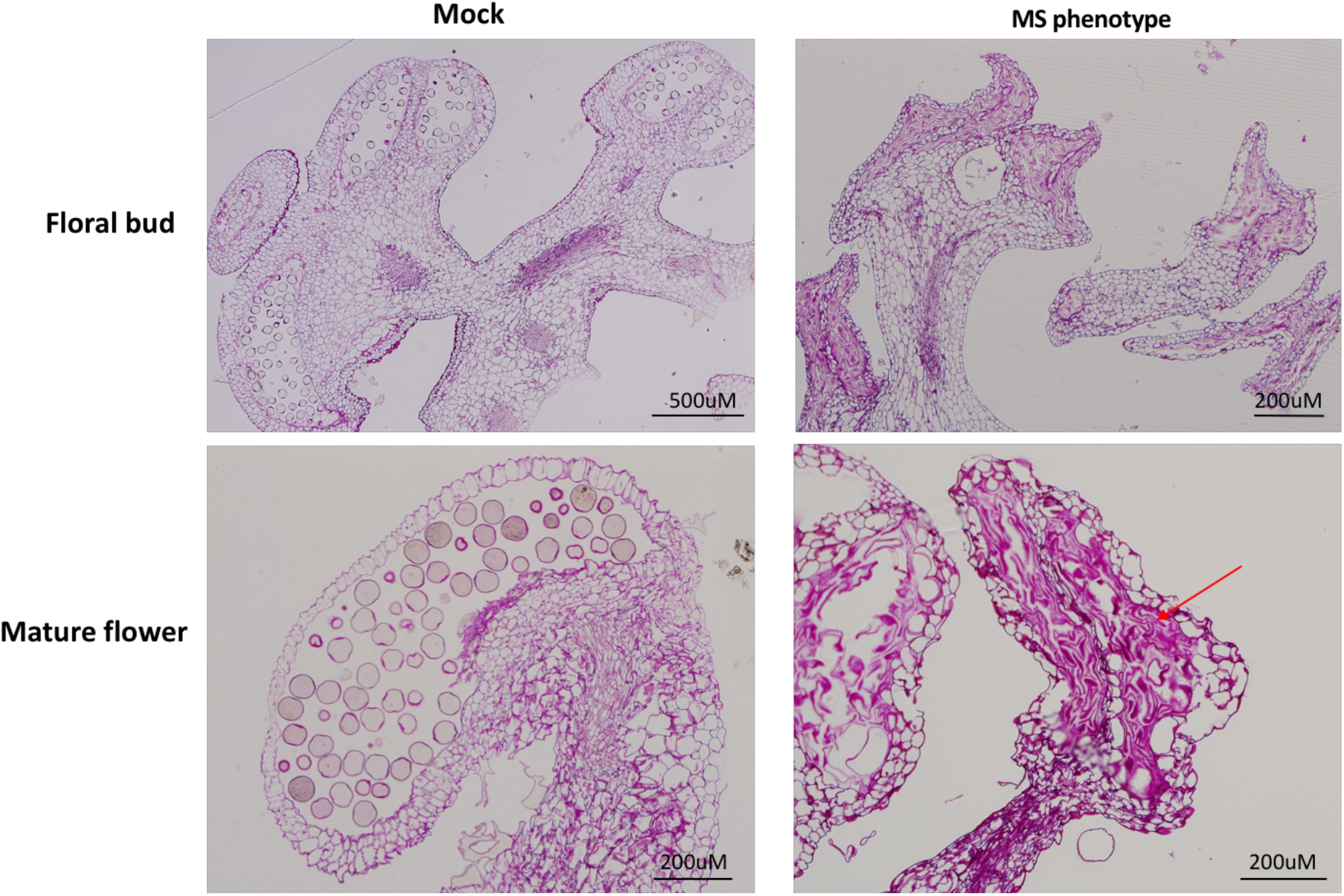
Histological sections of stamens from floral buds and mature flowers of mock-inoculated and candidate gene-silenced plants. The floral buds of plants infected with pCF93-pds-int (denoted as ‘mock’) formed normal pollen sacs containing pollen grains. In contrast, floral buds of candidate gene-silenced plant showed an absence of pollen. Pollen sacs of mature flowers exhibited normal morphology in mock flowers, but those of candidate gene-silenced plants were atrophied (red arrow). MS flower collected from an EXPA9-silenced plant (denoted as ‘MS phenotype’). Scale bars are 200 µm and 500 µm.

**Figure 6.**
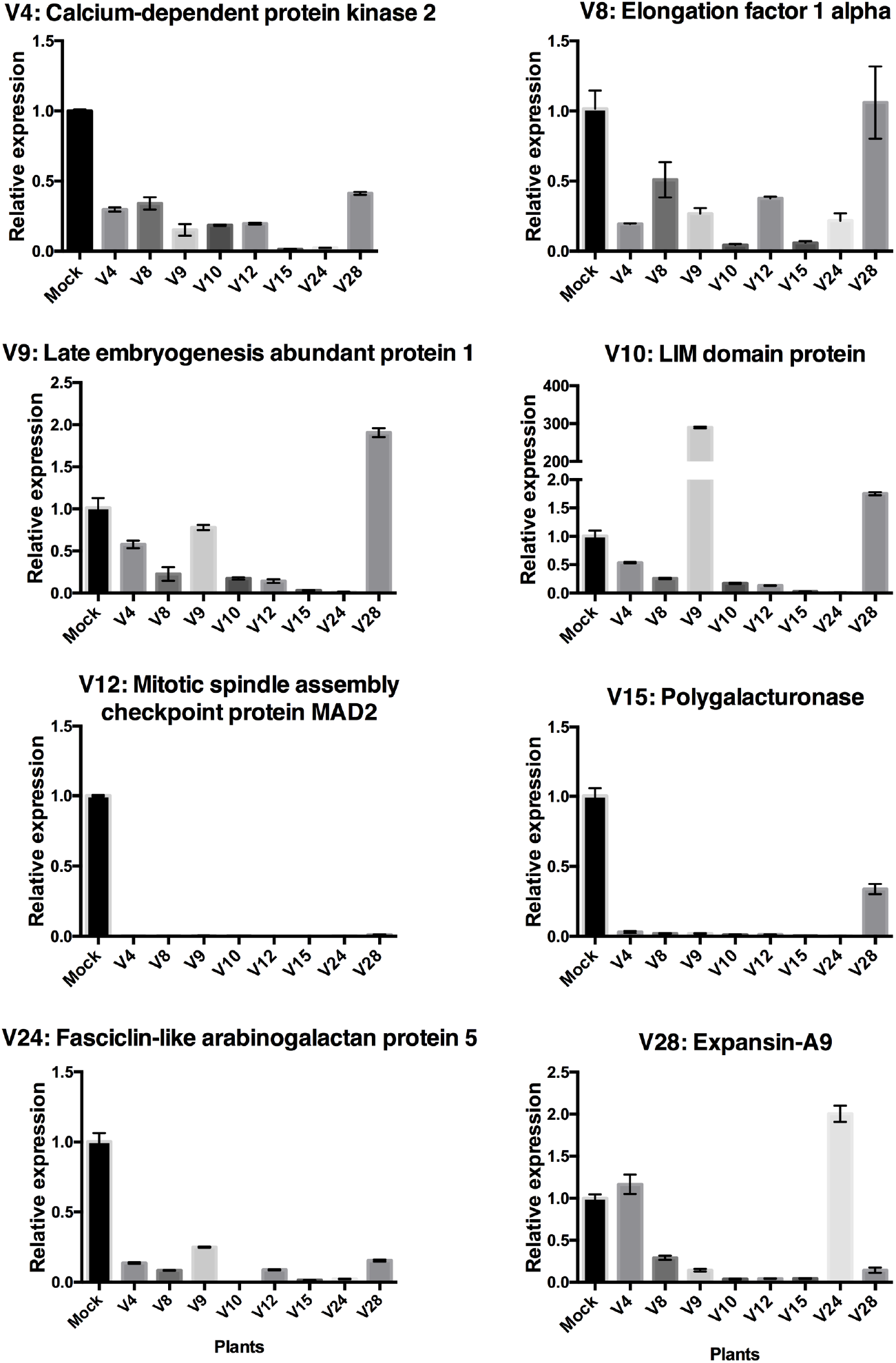
Male sterility-related candidate gene expression. Transcript levels were analyzed via RT-qPCR. Relative expression was normalized to 18S rRNA as a housekeeping gene. The transcript levels of eight candidate genes (Calcium-dependent protein kinase 2, Elongation factor 1 alpha, Late embryogenesis abundant protein 1, LIM domain protein, Mitotic spindle assembly checkpoint protein MAD2, Polygalacturonase, Fasciclin-like arabinogalactan protein 5, and Expansin-A9) were analyzed in all construct-infected plants. Each graph presents the relative expression of a candidate gene in the gene-silenced plants (x-axis). Relative expression levels were calculated by setting the expression level of pCF93-pds-int (denoted as ‘mock’) to 1.0 in arbitrary units (y-axis). Statistical analysis was performed using one-way ANOVA, followed by Tukey’s post hoc test for multiple comparisons.

## Discussion

Due to limitations associated with the genetic editing of cucurbits, genomic research into these species has been slower when compared to other plants. To address this issue, we previously isolated and constructed a CFMMV-Cm-based full-length clone (Rhee et al., 2014) and a virus vector for heterologous expression (Rhee et al., 2016). Fourteen CFMMV vectors were developed, carrying −55, −77, −81, −93, −100, −110, −121, −127, −148, −152, −157, −170, −180, and −187bp upstream positions from the CP start codon to the +100bp SGP, respectively. In previous studies on *Tobacco mosaic virus* (TMV), viral gene expression was reported to depend on the proximity of the ORF to the 3’ terminus (Culver et al., 1993). When the MP gene was located before the 3’ terminus, expression increased up to 20 times relative to that of the wild-type virus (Culver et al., 1993). The insertion of an additional 3’NTR between two coat proteins led to similar expression levels (Lindbo, 2007). Thus, it was suggested that proximity to the 3’NTR is a crucial factor for enhancing expression. TMV vectors were also designed with duplication of the TMV 3’ NTR between the foreign ORF and CP gene (Shivprasad et al., 1999). Based on these previous findings, two types of virus vectors were developed herein, one possessing a 3’NTR and the other possessing tandemly duplicated 3’NTRs, named CFMMV and CFMMV-K, respectively.

Among the previously developed vectors, −93 and −157 exhibited high EGFP expression efficiency. Therefore, pCF93, pCF93K, pCF157, and pCF157K were inoculated to evaluate VIGS efficiency in the current work. *N. benthamiana* infected with pCF93-nbpds, pCF93K-nbpds, and pCF157-nbpds developed photobleaching phenotypes, while gene silencing was unsuccessful in the pCF157K-nbpds-infected plant (Figure 1). pCF93K-nbpds and pCF157K-nbpds inserts were easily lost due to homologous recombination. *N. benthamiana* infected with pCF93-nbpds exhibited 5-fold lower levels of *PDS* mRNA in the leaves relative to pCF157-nbpds-infected plants.

pCF93-pds successfully induced gene silencing not only in leaves but also in the reproductive organs of cucurbits, including melon, cucumber, and watermelon. PDS is involved in the β- carotene biosynthesis pathway which is implicated in the color of flowers. Moreover, *PDS* downregulation can lead to inhibition of lycopene and β-carotene biosynthesis, resulting in the color change of watermelon flesh to white.

VIGS is sensitive to the genotypes of specific plant species (Bekele et al., 2019). Thus, three types of watermelon cultigens were evaluated in the current work. All cultigens exhibited gene silencing phenotypes, with fruits developing white-colored peels due to chlorophyll degradation and a reduction of lycopene content. Among evaluated cultigens, the DAH line showed the most pronounced silencing phenotypes and was selected for subsequent experiments. All three cultigens were small and could be grown in 16-cm pots with stable fruit development and ripening. Thus, our research required a relatively small cultivation area relative to common watermelon cultivation. Taken together, the VIGS approach described herein allows for convenient and relatively effortless watermelon research.

To functionally validate the 38 DEDEGs we identified, we inserted partial genes into the pCF93 vector which was then inoculated into DAH-MF seedlings via agroinoculation. Approximately 400 plants were inoculated and grown at once in the greenhouse (Supplemental Figure S1).Silencing eight of the 38 candidate genes seemed to induce male sterility, yielding different male sterile flowers within a single plant. To identify crucial genes implicated in male sterility, we evaluated the expression of these eight genes in MS flower samples. All genes exhibited a tendency toward downregulation in MS flowers, but no significant differences were observed in *EF1a, LEA1, LIM*, and *EXPA9* expression relative to mock-infected plants. EF1a is essential for translation, particularly elongation, and exhibits ubiquitous expression, which includes the reproductive organs of plants (Cosgrove et al., 1997; Wang et al., 2004). Thus, the downregulation of EF1a might affect male sterility-related gene expression. LIM, LEA, and Expansin play an important role in pollen formation and pollen tube elongation (Ye and Xu, 2012). Thus, we suspected that these four genes affect male sterility but might not be directly involved in the mechanism underlying MS. Of note, *CDPK2, FLA5, PG*, and *MAD2* were downregulated in all gene-silenced samples. There are several reports suggesting that CDPKs are involved in the regulation of pollen tube growth (Myers et al., 2009; Li et al., 2018). Thus, we suspect that CDPK2 downregulation might be indicated by the absence of pollen grain. PG is a representative hydrolytic enzyme involved in various functions such as plant cell wall deposition, fruit ripening, abscission, pollen intine and exine formation, as well as pollen tube growth via pectin rearrangement (Kulikauskas and McCormick, 1997; Huang et al., 2008; Jiang et al., 2014; Lyu et al., 2015). As our virus-infected MS flower did not produce pollen grain and exhibited an immature stamen, PG deficiency might have compromised stamen formation and pollen production. FLAs are subclass of arabinogalactan proteins (AGPs), which have pleiotropic functions implicated in plant reproduction and pollen development (Pennell and Roberts, 1990; Coimbra et al., 2007). AtFLA3 (*Arabidopsis thaliana FLA3*) knock-down plants produced abnormal pollens, and an overexpression line exhibited defective stamen filaments (Li et al., 2010). It was therefore, suggested that FLA3 participates in microspore and reproductive organ development.

Remarkably, *MAD2* expression was not detected in all virus-induced MS flowers (Figure 6). MAD2 is essential for chromosome segregation during mitosis, controlling the metaphase-to- anaphase transition (Shah and Cleveland, 2000). SAC is known as a crucial regulator for genome stability in humans, animals, and yeast (Lara-Gonzalez et al., 2012). While the functional significance of plant *MAD2* is still unclear, equal segregation of chromosomes is definitely essential for normal development. Arabidopsis and maize MAD2 homologues localized at the kinetochore, and maize MAD2 was reported to be sensitive to microtubule attachment during meiosis and mitosis (Yu et al., 1999). While the Arabidopsis MAD2-2 mutant did not affect reproductive organ development, it did compromise root growth. Lack of BUB1, which is one of the SAC components along with MAD2, induced male sterility in rice (Burgos-Rivera and Dawe, 2012). Taken together, the suppression of mitosis-related genes can give rise to male sterility in plants.

In the present study, we established a high-throughput screening system for the identification of putative male sterility-related genes deduced from massive RNA-seq data. This approach facilitated the functional genomic study of watermelon, which normally requires huge cultivation area in a short time for the simultaneous study of many genes of interest, in a common-sized greenhouse (Supplemental Figure S1). The use of small-fruit type watermelons planted in pots is, therefore, essential for reducing the study area, and it may also lower the risk of soil contamination.

The current work adds to the body of evidence regarding the major advantage of the VIGS system. The, pCF93 vector established silencing in the various organs throughout the lifetime of the plant. Taken together, pCF93 is a very useful tool for high-throughput screening and is particularly advantageous in fruit development.

To our knowledge, this is the first report on the application of VIGS for evaluating endogenous gene function in watermelon fruits, highlighting the utility of pCF93 in cucurbits. The current work establishes a theoretical and practical basis for future studies focused on functional genomics analysis in cucurbits.

## Materials and methods

### Plant materials

We used *Nicotiana benthamiana, Cucumis sativus* L. ‘Porisian pickling’, *Cucumis melo* L. ‘Early hanover’, as well as *Citrullus lanatus* L. ‘Chris cross’, ‘2401’, and ‘DAH’ for VIGS. The cultigens were obtained from the gene bank of the National Agrobiodiversity Center (NAC), Rural Development Administration (RDA), Jeonju, South Korea. The plants were grown under a constant temperature of 28°C with a 16/8-h light/dark cycle. Cucumber, melon, and watermelon plants were transferred to greenhouse 2∼3 weeks after inoculation. Each shelf in the green house is 150 × 270 cm and has 28 pots. For one experimental set of MS VIGS, we performed 14 replicates per putative gene. Twenty shelves were used for each set of MS VIGS experiments within the same time frame. Individual plants were planted in 16-cm pots and placed in the shelves at a distance of 15 cm from each other (Supplemental Figure S1).

### RNA extraction and cDNA synthesis

RNA was extracted using TRIZOL reagent (GIBCOBRL-Life technologies, MD, USA) and subsequently treated with 1 µl DNase (10 U/µl) at 37°C for 30 min. Two micrograms of total RNA were reverse-transcribed to cDNA by using Superscript III Reverse transcriptase (Invitrogen, CA, USA). The mixture included 2 µg of total RNA, 0.5mM of dNTP, and 100 ng of oligo dT or random hexamer and was incubated at 65°C for 5 min. Thereafter, the mixture was kept on ice for 1 min, followed by the addition of 5× first-strand buffer, 5mM DTT, 1 µl RNase inhibitor, 1 µl Superscript III Reverse transcriptase, and incubation at 50°C for 50 min as well as at 70°C for 15 min.

### Cloning of target insert genes for VIGS

PDS genes for VIGS were searched from GenBank. Fragments were amplified from *N. benthamiana PDS* (nbpds) and *Cucumis melo* L. *PDS* (cmpds) via RT-qPCR (Accession No. DQ469932 and KC507802, respectively). Nbpds was amplified by a primer set of Nbpds-F and Nbpds-R. Cucumber (cspds), melon (cmpds) were amplified by a primer set of cucurbit pds-F and cucurbit pds-R and watermelon *pds* (ccpds) was amplified by a primer set of ccpds-F and ccpds-R (Table S2). The *PDS* genes were cloned into pCF93, pCF157, pCF93K, and pCF157K (Figure 1). Putative male sterility-related genes were deduced from reference-based and *de novo* RNA-seq in male sterile and fertile watermelon lines previously reported by our group (Rhee et al., 2015; Rhee et al., 2017). pCF93-pds.int, constructed by inserting a partial fragment of pds intron into the MCS region of pCF93, was used as a mock. To amplify the intron region of watermelon pds gene, we isolated the genomic DNA from watermelon cultigen ‘Chris cross’ using DNeasy plant mini kit (Qiagen, CA, USA) followed the manufacturer’s manual. The target genes and primer sequences are presented in Table S1 and S2, respectively. The primer sets were designed based on the watermelon 97103 genome v1 at CuGenDB (http://cucurbitgenomics.org/).

Total RNA was reverse-transcribed to cDNA by using Superscript III Reverse transcriptase (Invitrogen, CA, USA) and oligo dT. cDNA was then amplified using Phusion DNA polymerase (Thermo, MA, USA). The PCR conditions were as follows: 98°C for 30 sec, 35 cycles of 98°C for 10 sec, 60°C for 30 sec, 72°C for 30 sec, and 72°C for 2 min. Each *Xho*I-*Pme*I-digested PCR fragment was ligated into the pCF93.

### Agroinoculation

All constructed vectors were transformed into *Agrobacterium tumefaciens* Gv3101. A single colony was picked up and cultured overnight in 5 ml LB media at 28°C. 100 µl of cultured cell-containing medium was transferred to fresh 30 ml LB medium containing 0.01M MES (pH 5.6) and 20 µM of acetosyringone. The cells were grown to an OD 600 of 1.0 and then harvested via centrifugation. Harvested cells were resuspended with MMA medium to a final O.D 600 of 0.9, and 200 µM acetosyringone was added. The inoculums were incubated at room temperature for 4 h with gentle agitation. The cells were inoculated onto cotyledons of cucurbits and *N. benthamiana*. Three true leaves per plant were used for inoculation. The agrobacterium solution was injected into the abaxial manner of leaves by using a 1-ml needleless syringe.

### RT-qPCR

We collected pooled leaf samples from individual *N. benthamiana* and cucurbit plants and isolated total RNA from three individual plants in triplicate in order to analyze VIGS efficiency via RT-qPCR. To determine gene expression levels in flowers, three individual flowers were used for total RNA isolation and cDNA synthesis. One microliter of cDNA was subjected to RT-qPCR. 18S rDNA and glyceraldehyde 3-phosphate dehydrogenase (GAPDH) were used as internal controls in *N. benthamiana* and cucurbits, respectively. RT-qPCR was conducted using Eco Real-Time PCR (Illumina, CA, USA). The PCR mixture included 10 µl 2× Realtime PCR mix (Biofact, Daejeon, Korea), 1 µl Evagreen dye, 10 µM primers, 1 µl cDNA, and ddH_2_ O up to a total volume of 20 µl. PCR conditions were as follows: 95°C for 12 min, 40 cycles of 95°C for 10 s, 60°C for 15 s, and 68°C for 15 s. We determined relative gene expression via the ddCT method. For statistical analysis of relative gene expression, one-way ANOVA Dunnett’s post hoc test was applied using PRISM 6 software.

### Quantification of chlorophyll contents

We collected 0.5 g of watermelon peels (approximately ten discs, 10 mm in diameter) and put them in 30 ml of absolute acetone overnight in the dark. Chlorophyll a and b contents were measured using a spectrophotometer and 1.00 cm quartz cuvettes with absorbance values at 651 and 664 nm, respectively. The formulae for calculating chlorophyll contents were as follows (Mackinney, 1941): chlorophyll a = (16.5 × O.D_664_ - 8.3 × O.D_651_) × A; chlorophyll b = (33.5 × O.D_651_ – 12.5 × O.D_664_) × A; total chlorophyll = (25.5 × O.D_651_ - 4 × O.D_664_) × A (A=Methanol volume (L) × 100πr^2^ (Cm) × number of leaf disc). A was calculated as 0.385 (A = (30/1000) × 100 × 3.14 × 0.5^2^x 10).

### Quantification of lycopene and β-carotene contents by using high-performance liquid chromatography (HPLC)

For sample preparation, 10 g of fresh watermelon flesh was freeze-dried for a week. The freeze-dried sample (∼ 0.1 g) was ground and mixed with silica beads (3-5 µm) (1:1, w/w). One milliliter of ethanol containing 0.5 mM BHT was added to the samples and struck with beads for 30 s. The bead-containing liquid samples were then transferred to 15-ml tubes, to each of which, 3 ml of petrol ether and 8 ml of 20% sodium chloride were gradually added with vortexing. The sample was centrifuged at 3000 rpm for 10 min, and the supernatant was collected. Finally, Na_2_ SO_4_ was added to the supernatant, which was then filtrated via a PTFE filter (13 mm, 0.2 µm; Advantec, USA). The pre-treated samples were analyzed through HPLC by using a Waters chromatography system equipped with reverse-phase column liquid (Kinetex 2.6 µm, C18 100a, 100 × 460 mm; Phenomenex, USA).

### Histological analysis and microscopy

To investigate the anther developmental differences between the MS and MF lines, floral buds and mature flowers were collected from both lines. The samples were soaked in 2.5% glutaraldehyde (v/v) for 90 min and washed in 0.1 M phosphate buffer (pH. 7.2). The samples were then soaked in 1% osmic acid (v/v) for 90 min and washed in 0.1 M phosphate buffer (pH. 7.2) at 4°C. Subsequently, the fixed samples were dehydrated using an ethanol gradient (40, 60, 80, 90, 95, and 100%) and then embedded in Epon-812 resin. Ultrathin cross-sections of 1500 nm thickness were prepared using an ultramicrotome (PT-X, RMC, USA) and stained with Periodic acid–Schiff (PAS) stain.

## Acknowledgements

We would like to show our gratitude to the gene bank of the National Agrobiodiversity Center (NAC) of Rural Development Administration (Gimje, Korea) for providing seeds. This work was carried out with the support of the “Cooperative Research Program for Agriculture Science & Technology Development (Project No. PJ01421302)’’ Rural Development Administration and the Golden Seed Project (213006055SBV20); the Ministry of Agriculture, Food, and Rural Affairs (MAFRA); the Ministry of Oceans and Fisheries (MOF); the Rural Development Administration (RDA); and the Korean Forest Service (KFS) of the Republic of Korea. This research was supported by the Chung-Ang University Graduate Research Scholarship in 2020.

## Data availability statement

All relevant data can be found within this manuscript and its supporting information files.

## Supplemental data

Supplemental Figure S1. Greenhouse plot layout to validate VIGS system for cucurbits.

Supplemental Figure S2. RT-PCR and sequence analysis of progeny virus from pCF157K-nbpds to confirm the homologous recombination in *N. benthamiana*.

Supplemental Figure S3. Male sterile flowers are produced in candidate gene silenced watermelon.

Supplemental Table S1. List of 38 de novo-exclusive differentially-expressed genes (DEDEGs) to validate gene function by VIGS.

Supplemental Table S2. Primers used for the construction of VIGS vectors.

Supplemental Table S3. Primers for RT-qPCR.

